# Proteomic Remodeling During Tumor Cell-Induced Platelet Aggregation Unveils Metastatic Drivers in Colorectal Cancer

**DOI:** 10.1101/2025.04.09.647951

**Authors:** Thorben Sauer, Caroline Gruner, Katharina Kern, Antje Rackisch, Lea Tischner, Katharina Schulz, Jasmin Ostermann, Lena Cohrs, Michael Kohl, Admar Verschoor, Timo Gemoll

## Abstract

**Background:** Colorectal cancer (CRC) is frequently associated with metastasis, resulting in high mortality rates. Platelets are known to play a crucial role in the metastatic cascade influencing tumor microenvironment remodeling, promoting cell transformation, facilitating metastatic niche formation, and shielding circulating tumor cells from immune surveillance. However, platelet proteomic alterations during tumor cell-induced platelet aggregation (TCIPA) remain largely unexplored. This study aims to characterize the proteomic profile of TCIPA in CRC using an in vitro model that recapitulates key aspects of CRC metastasis.

**Methods:** TCIPA was assessed via light transmission aggregometry using an *in vitro* model incorporating paired primary and metastatic cell cultures. Stable Isotope Labeling with Amino Acids in Cell culture (SILAC) allowed for the discrimination of healthy platelet and tumor cell proteomes prior to and following TCIPA. Data-independent acquisition mass spectrometry was employed to analyze intra- and extracellular tumor and platelet proteomes. Comparative proteomic profiling was performed using a range of bioinformatic analyses, including clustering, differential expression, and Gene Set Enrichment Analyses (GSEA).

**Results:** Comparison of the baseline proteome profiles of the CRC cell lines SW480 and SW620 identified 263 significant differentially expressed proteins (FDR ≤ 0.05, log_2_FC ≥ 1). The GSEA demonstrated enrichment of the ‘epithelial-mesenchymal transition’ (FDR: 5.617 × 10^−5^) gene set in SW480 cells. While SW480 exhibited rapid TCIPA, SW620 did not consistently interact with healthy platelets. Following TCIPA, 34 tumor proteins showed differential expression compared to their naïve status (without platelet-exposure). Notably, 17 of these proteins were significantly associated with CRC progression, particularly in the promotion of EMT, metastasis, tumor cell survival, proliferation, and metabolic reprogramming.

**Conclusions:** This study successfully characterized the proteomic profiles of platelets, platelet secretomes, and colorectal tumor cells following TCIPA-induced activation. The findings highlight the significant role of several tumor proteins and their metabolic effects in colorectal cancer progression, particularly with regard to metastasis.

## Background

In 2020, colorectal carcinoma (CRC) was responsible for over 1.9 million new cases and approximately 1 million deaths worldwide, solidifying its position as a leading cause of cancer-related morbidity and mortality (1). Notably, CRC is the second most commonly diagnosed cancer in women (11.5%) and the third in men (12.8%) (2). Considering an aging society, epidemiological forecasts suggest a substantial increase in the global CRC burden, with projections estimating 3.2 million new cases and 1.6 million deaths by 2040 (1). The development of metastatic disease presents a critical clinical challenge due to its association with increased mortality, mass effect, and disruption of physiological homeostasis. Approximately 20 – 25% of CRC patients present with metastatic spread at initial diagnosis, exhibiting a 5-year disease-free survival rate of 25%, emphasizing the imperative for innovative diagnostic and treatment interventions (3, 4).

In this context, the interaction of platelets with tumor cells (tumor cell-induced platelet aggregation, TCIPA) is recognized as a critical antecedent for successful metastatic dissemination. Platelets stabilize tumor cell arrest within the vasculature and protect tumor cells against the hostile environment in the bloodstream (5, 6). Evidence further indicates platelet migration into tumor tissue, where platelet-induced epithelial-mesenchymal transition (EMT) signaling serves as a key promoter of metastasis (7). Platelets exhibit dynamic responsiveness to pathophysiological conditions, including the capacity to uptake mRNAs and proteins from their surrounding environment (8–10). Consequently, the specific interaction with circulating tumor cells (CTCs) can modulate the protein composition in platelets, generating cancer-educated platelets (CEPs) that may subsequently exert additional pro-cancerous effects. Platelets possess the capacity to influence tumor growth and neovascularization through the release of transforming growth factor beta (TGF-β), vascular endothelial growth factor (VEGF), and platelet-derived growth factor (PDGF) (6). Furthermore, platelet-mediated processes, including the preparation of pre-metastatic niches through the modulation of granulocytes, the release of chemokines such as CXCL5 and CXCL7, and the release of matrix metalloproteases upon contact with tumor cells, have been demonstrated to be crucial for metastatic dissemination (11–13).

In this study, we employed stable isotope labeling with amino acids in cell culture (SILAC) coupled with data-independent acquisition mass spectrometry to investigate the interaction between healthy platelets with primary and metastatic CRC cell lines. Through differential expression and pathway enrichment analysis, we characterized functional alterations in the proteome of tumor cell-activated platelets, their corresponding secretome, and the activating cancer cells. Furthermore, we identified a panel of potential biomarkers associated with TCIPA-based CRC progression, potentially facilitating translation into clinical applications and improved patient outcomes.

## Material and Methods

### Cultivation of SW480 and SW620 cells

SW480 and SW620 cells were obtained from the American Type Culture Collection (ATCC, Manassas, VA, USA) and cultured as previously described (14). Specifically, cells were detached using Accumax^TM^ (PAN-Biotech GmbH, Aidenbach, Germany) and subsequently washed with PBS. The resulting cell pellet was resuspended in Tyrode’s buffer (140 mM sodium chloride, 3 mM potassium chloride, 16.62 mM sodium hydrogen carbonate, 1 mM magnesium chloride hexahydrate, 10 mM HEPES, 5.5 mM D-(+)-glucose, in MilliQ water, pH 7.4). The tumor cell suspension was adjusted to a final concentration of 5 × 10^5^ cells/mL.

### Stable isotope labeling with amino acids (SILAC)

To facilitate the discrimination of platelet-derived proteins from tumor cell proteins in the subsequent mass spectrometric analyses, stable isotope labeling by amino acids in cell culture (SILAC) was employed. Specifically, SW480 and SW620 cell lines were cultured in media supplemented with isotopically labeled arginine (13C15N-arg) and lysine (13C15N-lys). This SILAC strategy introduced a mass shift of 8 Da and 10 Da for lysine and arginine, respectively, enabling the differentiation between tumor cells (SILAC-labeled) and platelets (unlabeled). Culture labeling was performed using SILAC RPMI medium (SILANTES, Munich, Germany) supplemented with the labeled amino acids (arginine 200 mg/L, lysine 41.75 mg/L; SILANTES, Munich, Germany), 10 % dialyzed FBS, 5 mL glutamine (40 mg/L), and 1 % penicillin/streptomycin. Tumor cells were cultured in SILAC RPMI medium until an incorporation rate of 95% was achieved prior to aggregation experiments. For the determination of incorporation efficiency, 1 × 10^6^ cells were centrifuged, washed twice with PBS, and lysed in 100 μL of lysis solution and 1 µL of Nuclease, both components of the EasyPep Mini MS Sample Prep Kit (Thermo Fisher Scientific, Waltham, MA, USA). The resulting peptide lysates were subsequently analyzed by liquid chromatography-tandem mass spectrometry (LC-MS/MS) and the labeling efficiency was calculated using the MaxQuant (v1.6.1.0) software as described by Deng et al. (15).

### Isolation of platelets from whole blood

Platelets were isolated from plasma samples obtained from healthy volunteers. The Ethics Committee of the University of Lübeck gave ethical approval for this work (#19-147-A, vote of 15.04.2019). All patients provided informed written consent.

Upon receipt, the citrated blood tubes were immediately centrifuged at 150 × g for 20 min at room temperature without rotor brake. The platelet-rich plasma (PRP) was carefully transferred to a 15 mL reaction vessel using a Pasteur pipette. Premature aggregation of platelets was prevented by adding 400 nM PGI2 (prostacyclin). The PRP was then diluted 1:3 with Tyrode’s buffer and centrifuged at 800 × g for 10 min without rotor brake. The supernatant was completely removed and discarded. The sedimented platelets were resuspended in 500 μL platelet buffer, pooled, and adjusted to a concentration of 3 × 10^5^ platelets/μL to reflect the mean physiological platelet concentration in whole blood (16). The resulting platelet-rich suspension was maintained under gentle agitation at 10 rpm on a rocking shaker for 45 min.

### Light transmission aggregometry of TCIPA and sample collection

Light transmission aggregometry (LTA) measurements were performed using an APACT 4S Plus aggregometer (DiaSys Greiner GmbH, Flacht, Germany) with APACT LPC software AS-IS (v1.21c).

All samples were adjusted to a final concentration of 1 mM CaCl_2_ after 300 seconds (Carl Roth GmbH & Co. KG, Karlsruhe, Germany) to ensure consistent platelet aggregation (17). Aggregation was induced by the addition of either 12.5 μM TRAP-6 (Thrombin Receptor Activator Peptide 6) or 20 μL of tumor cell suspension (2,500 cells/μL) containing isotopically labeled SW480 or SW620 cells after baseline reading of 500 seconds. TRAP-6 activates platelets by selectively targeting the preotease-activated receptor 1 (PAR-1) on their surface. Aliquots of platelets alone, tumor cells alone, and the platelet/tumor cell mixture following TCIPA were collected. Platelets incubated with CaCl_2_ and TRAP-6 served as negative and positive controls, respectively. Immediately following collection, samples were centrifuged for 15 seconds and lysed using the EasyPep Mini MS Sample Prep Kit (Thermo Fisher Scientific, Waltham, MA, USA) before being stored at −80 °C until further analysis.

### Immunofluorescence microscopy

The aggregation assay was carried out on microscopy slides allowing immunofluorescence imaging of platelet-tumor interaction. Platelets were isolated from a single healthy donor. For each experimental condition (10-min and 23-min incubation), 1.5 µL of 200 mM CaCl_2_ was added to 300 µL of platelet solution (3 × 10^5^ platelets/µL) in 2 mL protein low-binding tubes. After 500 seconds incubation, 30 µL tumor cell solution (7.5 × 10^4^ cells) was added. Samples were incubated for 2 or 15 minutes, diluted 1:2 with 300 µL of Tyrode’s buffer, and centrifuged six times at 800 × g for 3 minutes with low acceleration. For the platelet-only protocol, 300 µL platelets were incubated with 1.5 µL CaCl_2_ and 300 µL of Tyrode’s buffer for 15 min and subsequently processed using cytospin preparation.

For immunofluorescence microscopy, cytospin preparations were air-dried overnight and fixed with ice-cold acetone for 15 minutes. After fixation, slides were washed for 1 minute on a shaker with PBS (1x, pH 7.4; Gibco, Thermo Scientific, Waltham, USA, CAT# A1286301). Nonspecific binding was blocked using 1% bovine serum albumin (BSA, Miltenyi, Bergisch Gladbach, Germany) in PBS for 30 minutes in a humidified chamber. Primary antibody staining was performed using anti-CD42b (Clone 42CO1, Invitrogen, Thermo Scientific, Waltham, MA, USA, CAT# MA5-11642) diluted 1:100 in 0.1% BSA/PBS. 100 µL of antibody solution was applied to each cytospin and incubated for 1 hour at room temperature in a humidified chamber. The slides were then washed three times with PBS for 5 minutes each. Secondary antibody staining was conducted using Alexa Fluor 488-conjugated anti-mouse IgG (Abcam, Cambridge, UK, CAT# 150105) diluted in 1:500 in 0.1% BSA/PBS. 100 µL of the secondary antibody solution was applied to each cytospin and incubated for 1 hour at room temperature in a humidified chamber. Subsequently, the slides were washed three times with PBS for 5 minutes each and mounted using DAPI (4’,6-Diamidin-2-phenylindol) Fluoromount-G (SouthernBiotech, CAT# 0100-20). CD42b expression and nuclear staining were visualized using a Keyence BZ-9000 (Osaka, Japan).

### Sample preparation for mass spectrometry

Protein concentration of the cell lysates was determined using the EZQ Protein Quantitation Kit (Invitrogen, Waltham, MA, USA) according to the manufacturer’s protocol. Sample preparation was performed with the EasyPep Mini MS Sample Prep Kit (Thermo Fisher Scientific, Waltham, MA, USA) adhering to the manufacturer’s instructions. For each sample, 25 to 100 μg of protein was digested with the provided trypsin/Lys-C protease mix. Subsequently, the peptides were purified using purification columns. The samples were subsequently frozen at −80 °C and lyophilized for 1 hour 20 minutes using a Christ Alpha vacuum centrifuge. Lyophilized samples were stored at −20 °C until reconstitution for downstream analysis.

### Mass spectrometry profiling

Samples were solubilized to a final concentration of 1 µg/ml in solvent A (1 % formic acid (v/v) in HPLC/MS ultrapure water) and loaded into a Dionex Ultimate 3000 HPLC system (Thermo Fisher Scientific, Waltham, MA, USA). The samples were first loaded onto a trap-column (μ-precolumn Acclaim PepMap 100 C18, diameter × length: 0.3 mm × 5 mm, particle size: 5 μm, pore size: 100 Å, Thermo Fisher Scientific, Waltham, MA, USA) and desalted with solvent A at a flow rate of 5 µL/min for 4 min. Subsequently, the samples were separated using an analytical column (Luna C18(2): 0.3 mm x 50 mm, particle size: 3 μm, pore size: 100 Å, Phenomenex Inc., Torrance, CA, USA) and eluted with a multistep gradient of solution B (0.1 % formic acid (v/v) in acetonitrile) in solution A for 86 min at 5 µL/min.

The purified and separated peptides were analyzed using a TripleTOF 5600+ mass spectrometer (AB Sciex, Framingham, MA, USA) using electrospray ionization (ESI). The analysis of TCIPA samples was performed with data-independent SWATH-MS acquisition (sequential window acquisition of all theoretical mass spectra), using the following parameters: ion spray voltage, 5,000 V; ion source gas 1, 15; ion source gas 2, 0; curtain gas, 30; and source temperature heating set to 0°C; declustering potential, 100; collision energy, 25.1; collision energy spread, 5; ion release delay, 67; ion release width, 25. One MS1 scan was performed with an accumulation time of 50 ms in the mass-to-charge range of 350-1,250 m/z, followed by 100 Q1 windows with variable m/z width ranging from 5 to 85.6 Da, each at an accumulation time of 30 ms and a collision energy spread of 5 eV. The precursor isolation windows were defined using the SWATH variable window calculator (V1.1, AB Sciex, Framingham, MA, USA) based on precursor m/z densities obtained from DDA spectra.

DDA-MS measurements were performed to generate the SWATH method and to assess the incorporation rate of the isotopically labeled amino acids in the SILAC approach. Identical instrument parameters as stated above were used. An MS1 scan was performed for 350-1,250 Da with an accumulation time of 250 ms followed by MS2 scans for 100-1,500 Da with an accumulation time of 50 ms at high sensitivity mode.

### Incorporation efficiency of isotope-labeled amino acids

To ensure complete incorporation of the isotope-labeled amino acids into tumor cells prior to aggregation experiments, incorporation efficiency was determined. Cellular proteins of labeled SW480 and SW620 cells were analyzed by LC-MS/MS using DDA. Protein inference was performed using MaxQuant software (v1.6.1.0, Max Planck Institute of Biochemistry, Berlin, Germany) and the human UniProtKB Swiss-Prot protein sequence database (18). Data analysis used a false discovery rate (FDR) of 0.01, excluding reverse proteins and contaminants. Methionine oxidation and N-terminal acetylation were included as variable modifications. Arg10 and Lys8 were selected as labels with trypsin and Lys-C as the digestive enzymes (19).

For peptides terminating in lysine or arginine, incorporation efficiency was calculated using ‘Intensity H’ and ‘Intensity L’, representing the summed extracted ion chromatogram of the isotopic cluster of labeled and unlabeled peptide components, respectively (15). Mean incorporation efficiency was determined separately for lysine- and arginine-ending peptides. Complete incorporation was defined as a mean incorporation efficiency of >95% (20).

### Identification and quantification of proteins

SWATH®-MS raw data of the platelet and tumor cell samples were processed using the multiplex functionality DIA-NN V1.8.1 software (data-independent acquisition by neural networks) (21, 22). Independent analysis runs were performed to generate distinct protein lists for platelets and tumor cells. An *in-silico* spectral library was predicted from the human UniProtKB Swiss-Prot database FASTA (version 202/12/6) (18) file using DIA-NN options for FASTA digest and deep learning-based prediction of spectra, retention times, and ion mobilities. Analysis of platelets and tumor cell proteins from raw MS/MS spectra was conducted primarily based on the default settings: Trypsin/P was specified as protease with a maximum of one missed cleavage. The maximum number of variable modifications was set to zero. N-terminal methionine excision and C-carbamidomethylation were enabled, whereas methionine oxidation, N-terminal acetylation, phosphorylation, and GG adduction on lysins were disabled and therefore not considered as variable modifications. The default settings were kept for the peptide length range (7-30 amino acids) and the precursor charge range (1–4). The precursor m/z range was set to 300-1,800 Da whereas the fragment ion m/z range was defined as 350-1,250 Da. The MS1 and MS2 accuracies were fixed to 15 and 20 Da, respectively. The ‘Match between runs’ and ‘No shared spectra’ options were enabled. Protein inference was performed based on genes, using single-pass mode as neural network classifier, and ‘Any LĆ (high accuracy) quantification strategy. The cross-run normalization was RT-dependent. For tumor cell protein identification, SILAC labeling was accounted for using additional command line options: ‘--fixed-mod Lys8,8.014199,K,label’; ‘--fixed-mod Arg10,10.008269,R,label’; ‘--lib-fixed-mod Lys8’; ‘--lib-fixed-mod Arg10’; ‘--original-mods’ and ‘--gen-spec-lib’. The DIA-NN output was further processed using the *DIA-NN R* package (v1.0.1) (21). Filtering criteria included precursor with library q-value and library protein group q-value ≤ 0.01. The dataset was filtered for proteotypic proteins, and the MaxLFQ algorithm was used for protein quantification (23). Proteins identified with less than two proteotypic peptides were excluded. The remaining proteins were finally mapped to their corresponding gene names for downstream analysis. The mass spectrometry proteomics data have been deposited to the ProteomeXchange Consortium (24) via the PRIDE partner repository (25) with the dataset identifier PXD062043.

The R package *DEP* (v1.22.0) was used for quantitative data preprocessing (26). Missing values were defined as absent data in the data matrix. The protein data matrices of the platelet proteomes (sediment and secretome) were filtered to include only proteins identified in all replicates of at least one condition/group. The tumor cell sediment dataset was filtered for full profiles due to large identification heterogeneity. Quantile normalization, including a log_2_ transformation, was performed on all datasets using the *QFeatures* package (v1.10.0). Normalization success was verified using MeanSD and RLE plots. For platelet proteome datasets, missing value imputation utilized multiple strategies based on the nature of the missingness. Proteins exhibiting missing values in all replicates of at least one condition were considered missing not at random (MNAR) and were imputed using random draws from a Gaussian distribution centered around a minimal value (27). In contrast, other missing values were classified as missing at random (MAR) and were imputed using the k-nearest neighbor algorithm. The R package *DEP* borrows the imputation function from MSnbase and was used with default settings (28, 29).

### Differential protein expression analysis

The semi-quantitative data of the two cell lines SW480 and SW620 were compared using the Limma method from the *limma* R package (v3.56.2) (30). Benjamini–Hochberg procedure (31) was employed for correction for multiple testing. Differential expression of proteins was considered significant at q ≤ 0.05 and at absolute logarithmic fold change (|log_2_FC|) of ≥ 1.

To identify differentially expressed platelet proteins involved in TCIPA, the protein levels of platelet-naïve (but recalcified) platelets were compared to those after interaction with SW480 or SW620 cells. As a positive control, the protein levels of platelets before and after activation with TRAP-6 also analyzed. To identify differentially expressed tumor proteins post-TCIPA, the protein levels of SW480 or SW620 tumor cells were compared to the control condition of platelet-naïve (but recalcified) SW480 or SW620 cells.

To compare the protein expression of activated platelets and tumor cells to their corresponding controls, a Welch analysis of variance (ANOVA) was computed for each dataset using the *stats* R package (v4.3.1) for differential protein expression analysis. The results of the Welch ANOVA were corrected for multiple testing using the Benjamini–Hochberg procedure. Post-hoc testing was performed via the Tukey honestly significant difference (HSD) method (32) utilizing the Tukey HSD function from the *stats* R package. Differential protein expression was defined significant at a q-value of ≤ 0.05 for both, ANOVA and post-hoc tests, along with an |log_2_ FC| of ≥ 1. Volcano plots were generated from the results of the differential expression analysis using the *ggplot* R package (v3.4.4) (33).

### Bioinformatic analysis

A principal component analysis (PCA) was computed using the base R *stats* package and visualized using the autoplot function of the *ggplot* package. A gene set enrichment analysis was performed using the *GSEA* function from the *ClusterProfiler* R package (4.8.3) (34) to functionally annotate the observed differences between the two colorectal cancer cell lines. The protein profiles were analyzed using the Gene Ontology database (35, 36) and the Molecular Signatures Database v7.4 (MSigDB) (37, 38), specifically the ‘Hallmark collection’ (39). Benjamini-Hochberg procedure was employed as a correction method for multiple testing, and significant enrichment was considered at a q-value ≤ 0.05. Additionally, target proteins characterizing the proteomic profile differences of platelet-exposed SW480 and SW620 were functionally analyzed with an overrepresentation analysis (ORA) using the *enricher* function of the *ClusterProfiler* R package and the MSigDB C2 collection (curated gene sets). Again, the Benjamini-Hochberg procedure was applied for multiple testing corrections, with significant overrepresentation defined at a q-value ≤ 0.05. Significant results were filtered to gene sets whose descriptions contain the terms ‘cancer’ and/or ‘metastasis’ and/or ‘metastatic’.

To visualize set intersections of differentially expressed proteins among different condition comparisons, Venn diagrams were created using the R package *ggvenn* (v0.1.10). A protein rank plot was created by calculating the average intensity across both conditions, highlighting the target proteins by coloring them based on their log_2_ FC derived from the differential expression analysis. The PubMed database was searched in order to identify relevant studies based on the corresponding Gene Symbol and ‘colon cancer’ and/or ‘colorectal cancer’. Each search was limited to studies published in English.

## Results

### The proteome profiles of SW480 and SW620 widely differ from one another

To establish a foundational reference for downstream experiments investigating the interaction between tumor cells and platelets, we characterized the proteomes of the CRC cell line pair SW480 (primary tumor-derived) and SW620 (lymph node metastasis-derived) under platelet-naïve conditions. This dataset encompasses intensity measurements for 3,027 proteins. Principal component analysis demonstrated distinct separation between the proteomic profiles of the two cell lines (Figure 1A). Differential expression analysis identified 263 proteins differentially expressed (Figure 1B, Supplementary Table S1, Additional File 1). Gene set enrichment analysis using the gene ontology (GO) sub-ontology ‘biological process’ revealed significant enrichment of 44 GO terms (FDR ≤ 0.05) in SW480 compared to SW620. Notably, significant enrichment of the adhesion-associated GO terms was observed (Supplementary Table S2, Additional File 1). Employing GSEA on the Molecular Signatures Database Hallmark gene set collection (39) revealed an association of the SW480 cells with the gene sets ‘Epithelial mesenchymal transition’, ‘Coagulation’, ‘IL2-STAT5 signaling’, and ‘Apoptosis’. Conversely, the ‘MYC Targets V2’ gene set was significantly enriched in SW620 (Figure 1C and Supplementary Table S2, Additional File 1). These results indicated distinct molecular signatures associated with each cell line, demonstrating functional differences correlated with their respective metastatic phenotype.

**Figure 1:**
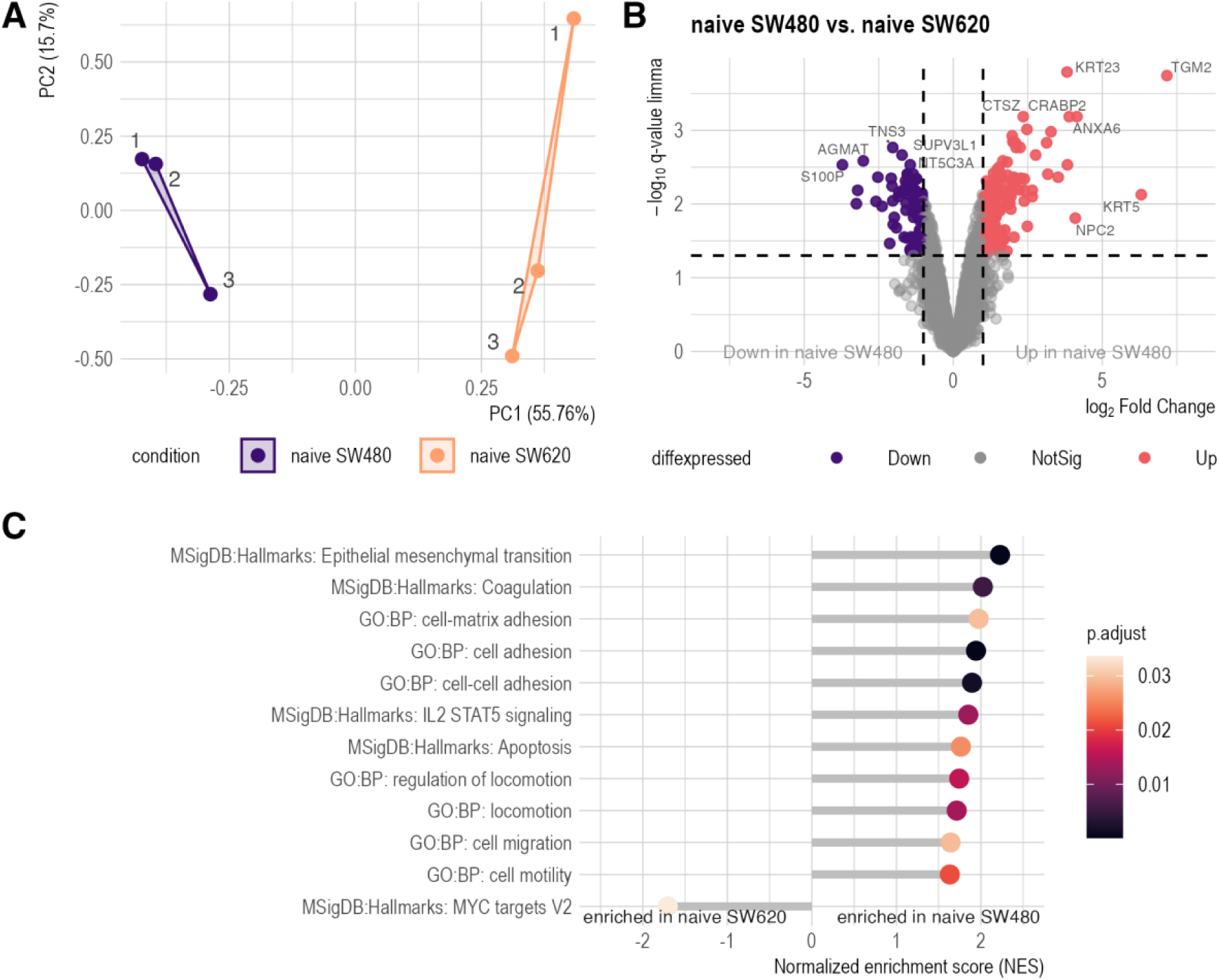
Proteomic characterization of SW480 and SW620. **A:** Principal component analysis (PCA) plot of 3,027 proteins, shows distinct clustering of naive SW480 and naive SW620 cell lines. Principle component 1 (PC1, x-axis) accounts for 57.76% of the variance, whereas PC2 (y-axis) explains 15.7%. **B:** Volcano plot illustrating differential protein expression between naive SW480 and SW620 cell lines. Significantly differentially expressed proteins (q ≤ 0.05 and |log_2_FC| ≥1) are highlighted, with 241 proteins meeting these criteria. Red points indicate proteins with significantly increased abundance, while purple points represent significantly decreased abundance of proteins. Differential expression analysis was conducted using limma. **C:** Selected gene set enrichment (GSEA) results using gene ontology (GO) Biological Processes subdatabase and the Molecular Signature Database (MSigDB) Hallmark collection. Enrichment was considered significant if q-value ≤ 0.05.

### Platelets from healthy donors are rapidly activated by TRAP-6 & SW480, but not by SW620

Four independent aggregation assays were performed for the tumor cell lines SW480 and SW620, with TRAP-6 serving as a positive control and CaCl_2_ as a negative control. While SW480 (Figure 2A) and TRAP-6 (Figure 2C) led to reproducible aggregation within 28 ± 3 minutes and 15 minutes, respectively, SW620 cells stimulated platelet aggregation in only two cases, with partial aggregation occurring after approximately 85 and 115 minutes (Figure 2B). The addition of CaCl_2_ (Figure 2D) did not trigger any aggregation throughout the duration of the experiments. Aggregation initiated by SW480 cells (reaching 20% aggregation) reached a high aggregation extent (80%) significantly later than that observed with TRAP. Consequently, SW480 cells exhibited a significantly prolonged aggregation interval (Figure 2E). While SW620-induced aggregation reactions exhibited delayed initiation and completion, the limited sample size precluded statistical evaluation (Figure 2E). In a separate experiment, immunofluorescence microscopy was utilized to capture the aggregation process on a glass slide. The aggregation reaction was terminated after ten minutes of aggregation to capture the initial state for both the SW480 and SW620 cells, which had been incubated with platelets, respectively (Figure 2F). Both initial conditions displayed a uniform distribution of platelets in a loosely organized formation (green staining) and widely dispersed cancer cells (blue staining). Following a 23-minute incubation, platelets incubated with SW480 showed extensive aggregation and cluster formation, partially surrounding the cancer cells (Figure 2G). In contrast, platelets exposed to SW620 exhibit initial signs of aggregation onset and early formation of platelet aggregates, but to a lesser degree compared to SW480. Recalcified platelets (CaCl2 condition) showed no evidence of aggregation after 23 minutes (Supplementary Figure 1, Additional File 2). Collectively, these results demonstrate that SW480 cells induce more rapid and consistent platelet aggregation compared to SW620 cells, supported by both light transmission aggregometry and immunofluorescence microscopy.

**Figure 2:**
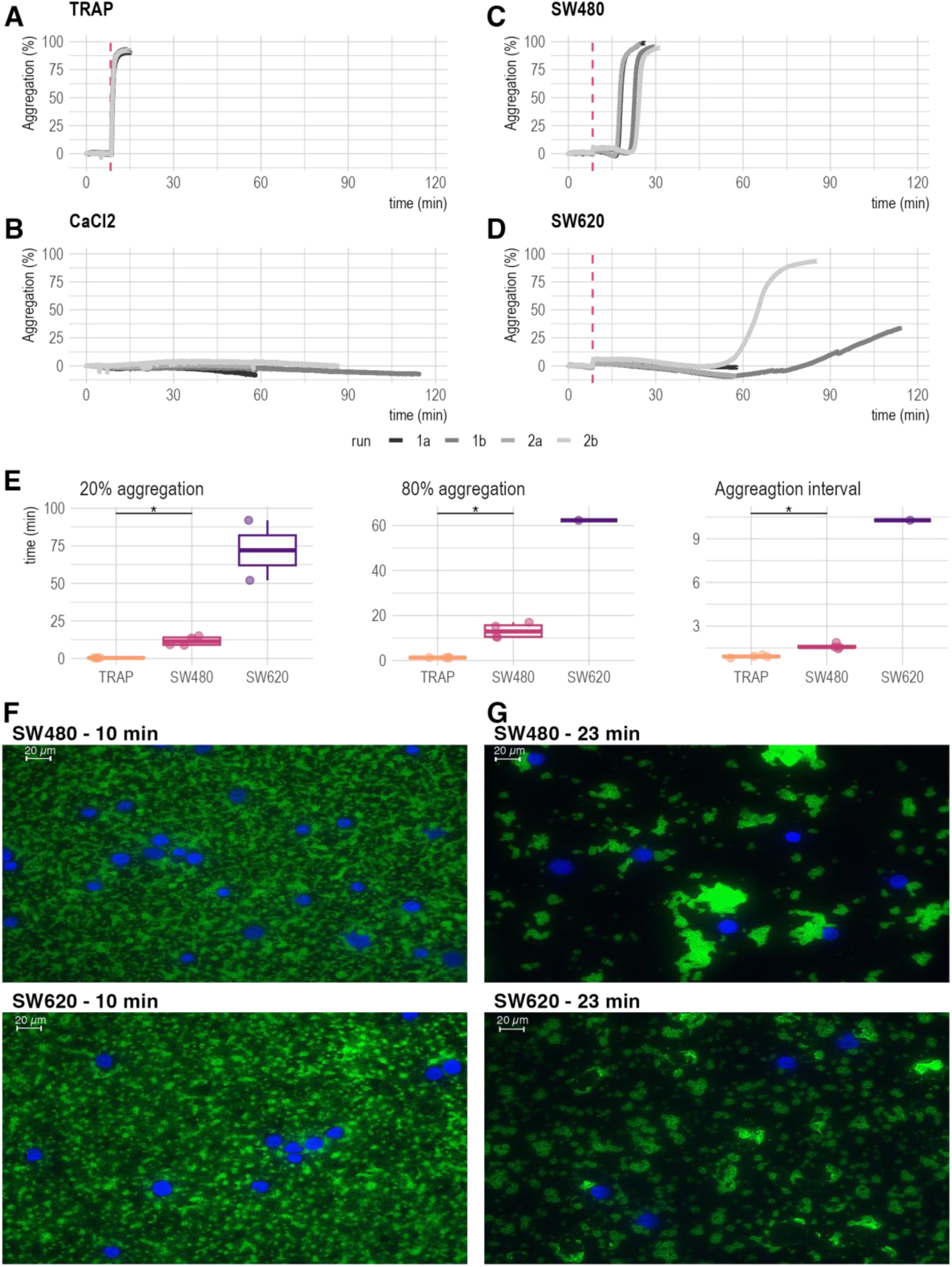
Light transmission aggregometry (LTA) and Immunofluorescence Analysis of TCIPA. **A-D:** LTA profiles demonstrating distinct platelet aggregation patterns, dashed red line indicates agonist addition: **A:** TRAP-6 (positive control) eliciting the fastest and most consisting platelet aggregation. **B:** CaCl_2_ (negative control) showing no platelet aggregation over up to 115 minutes. **C:** SW480-induced rapid and reproducible platelet aggregation. **D:** SW620-induced delayed and inconsistent platelet aggregation (only 2 out of 4 samples, >60 min). **E:** Boxplot summary of aggregation times. Time (in minutes) to reach 20% and 80% aggregation is shown for samples achieving these thresholds. SW620 samples exhibit the longest and most variable aggregation times, with the fewest samples reaching the thresholds. Statistical significance was determined by the Wilcoxon rank sum test (p ≤ 0.05). **F:** Immunofluorescence microscopy of TCIPA at 10 and 23 minutes for SW480 and SW620 cell lines (40x magnification). Cell nuclei stained with DAPI (blue) and platelets labeled with anti-CD42b antibody (green). At 10 minutes, no aggregation is visible in either cell line. At 23 minutes, SW480 cells show clear platelet aggregation and cluster formation around tumor cells, while SW620 cells exhibit reduced platelet aggregation and cluster formation.

### The platelet protein profile and secretion changes upon aggregation

LC-MS/MS analysis of platelet proteomes identified 2,379 unique proteins, corroborating the TCIPA profile on the proteomic level. A PCA clearly separated TRAP-6 and SW480 conditions from the CaCl_2_ and SW620 experiments (Figure 3A). Notably, partially aggregated SW620 samples (1b & 2b) clustered distinctly from the negative control (CaCl_2_) and the non-aggregated SW620 samples. Differential abundance calculation (ANOVA adj. p-value & post hoc q-value ≤ 0.05, and |log_2_FC| ≥ 1) revealed 24 significantly altered proteins in the SW480 *versus* CaCl_2_ comparison (Figure 3B, Supplementary Table S3, Additional File 1). Comparison of SW620 *versus* TRAP-6 identified 27 proteins (Supplementary Figure 2C, Additional File 2; Supplementary Table S3, Additional File 1), while only three proteins were identified in the SW620 *versus* CaCl_2_ comparison (Figure 3C, Supplementary Table S3, Additional File 1). In line, the comparison of SW480 and SW620 exposed platelet proteomes identified 21 proteins as being highly differentially abundant (Figure 3D Supplementary Table S3, Additional File 1). The differential expression analysis results of all other comparisons are visualized in supplementary Figure 2 and listed in Supplementary Table S3, Additional File 1. Collectively, these findings demonstrate distinct interaction characteristics of SW480 and SW620 with healthy platelets, allowing for categorization of proteins into negative and positive platelet aggregation patterns.

**Figure 3:**
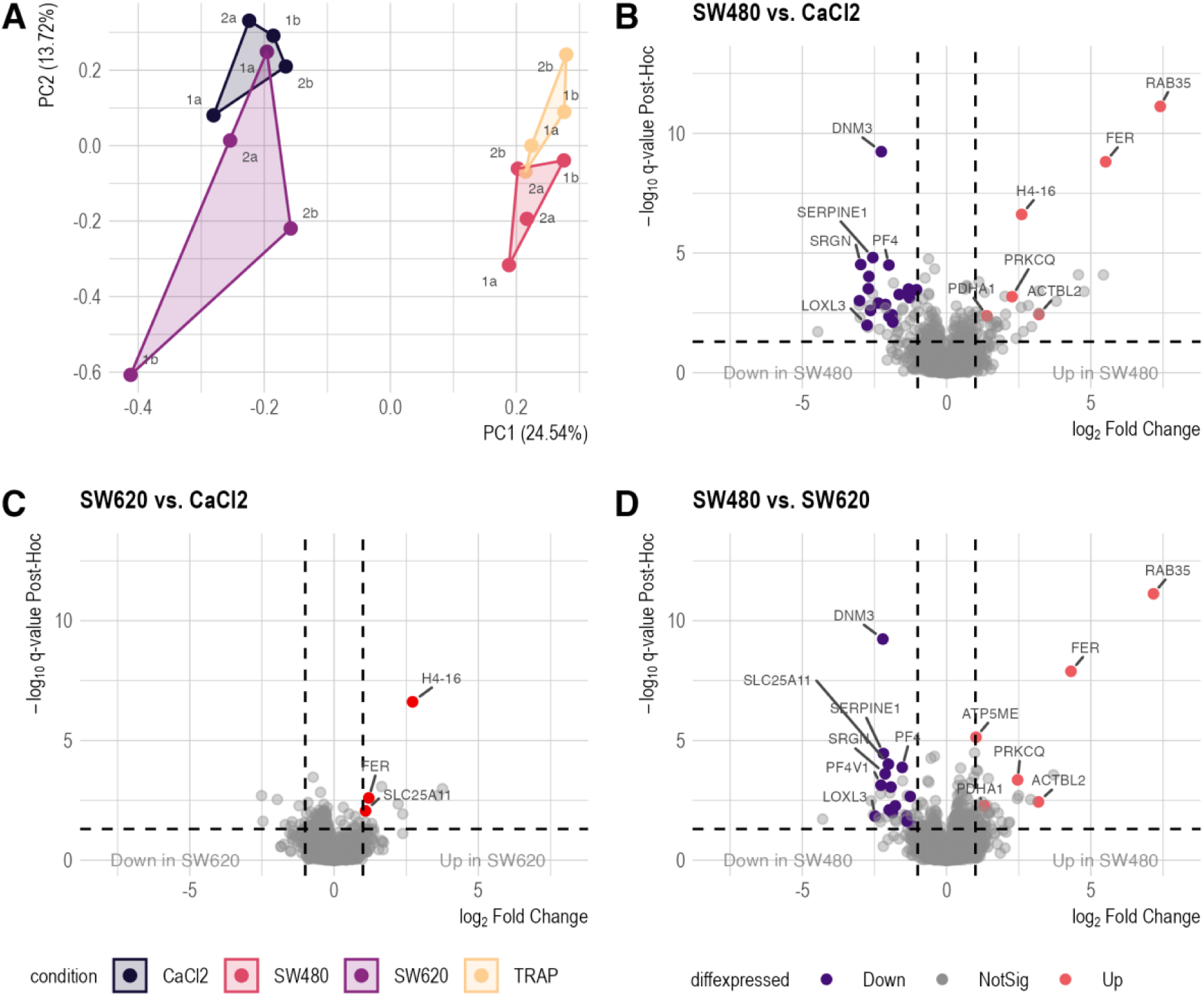
Proteomic analysis of platelet aggregation-induced changes. **A:** Principal component analysis of 2,379 proteins reveals distinct clustering patterns across different aggregation conditions. TRAP-6 and SW480-induced samples are clearly separated from the CaCl_2_ negative control, with SW620-induced samples showing a less pronounced differentiation. Notably, TRAP-6 and SW480 samples exhibit remarkable proximity in the PCA plot, suggesting similar proteomic responses. PC1 accounts for 24.54% of the variance, while PC2 explains 13.72%. **B-D:** Volcano plots visualize results of differential expression analysis using ANOVA and Tukey HSD post-hoc testing, comparing different aggregation conditions. Proteins are considered significantly differentially expressed at an ANOVA and post-hoc q ≤ 0.05, and |log_2_FC| of ≥1). Red points indicate proteins with significantly increased abundance, while purple points represent significantly decreased abundance of proteins.

Subsequently, an analogous analysis was performed on the secreted platelet proteome. Based on 546 unique proteins identified, PCA clustering (Figure 4A, Supplementary Table S4, Additional File 1) demonstrated a clustering pattern comparable to the behavior observed for the cellular platelet sediment (Figure 3A). A protein-wise comparison of the different experimental conditions to the negative control revealed significant differences in the proteomic profiles of the TRAP-6 and SW480 conditions, but not the SW620 (Supplementary Figure 3A-C, Additional File 2; Supplementary Table S4, Additional File 1). In contrast, only one protein was identified as differentially abundant in the SW620 *versus* CaCl_2_ comparison, suggesting that SW620, the metastatic counterpart of the SW480 cell line, did not evoke a proteomic change of the platelet secretome (Supplementary Figure 3C, Additional File 2, Supplementary Table S4, Additional File 1). The proteomic profiles of SW480-stimulated and TRAP-activated platelet proteomes significantly differed, with 52 differentially expressed proteins identified (Figure 4B, Supplementary Table S4, Additional File 1). Analysis of SW480-stimulated platelet secretomes identified four proteins with uniquely altered abundance compared to the positive control, suggesting specific effects of SW480 cells on platelet activation (Figure 4C). The abundance of fibrinogen alpha, beta, and gamma (FGA, FGB, and FGG) was significantly decreased in the SW480-stimulated platelet secretomes, whereas Glycoprotein 5 Platelet (GP5) showed increased levels (Figure 4D). These results demonstrate distinct interaction patterns between primary (SW480) and metastatic (SW620) CRC cells: SW480 cells induce significant changes in platelet protein profiles and secretion, while SW620 cells show minimal impact, suggesting that different stages of cancer progression uniquely influence platelet function and potentially affect tumor-platelet interactions during the metastatic process.

**Figure 4:**
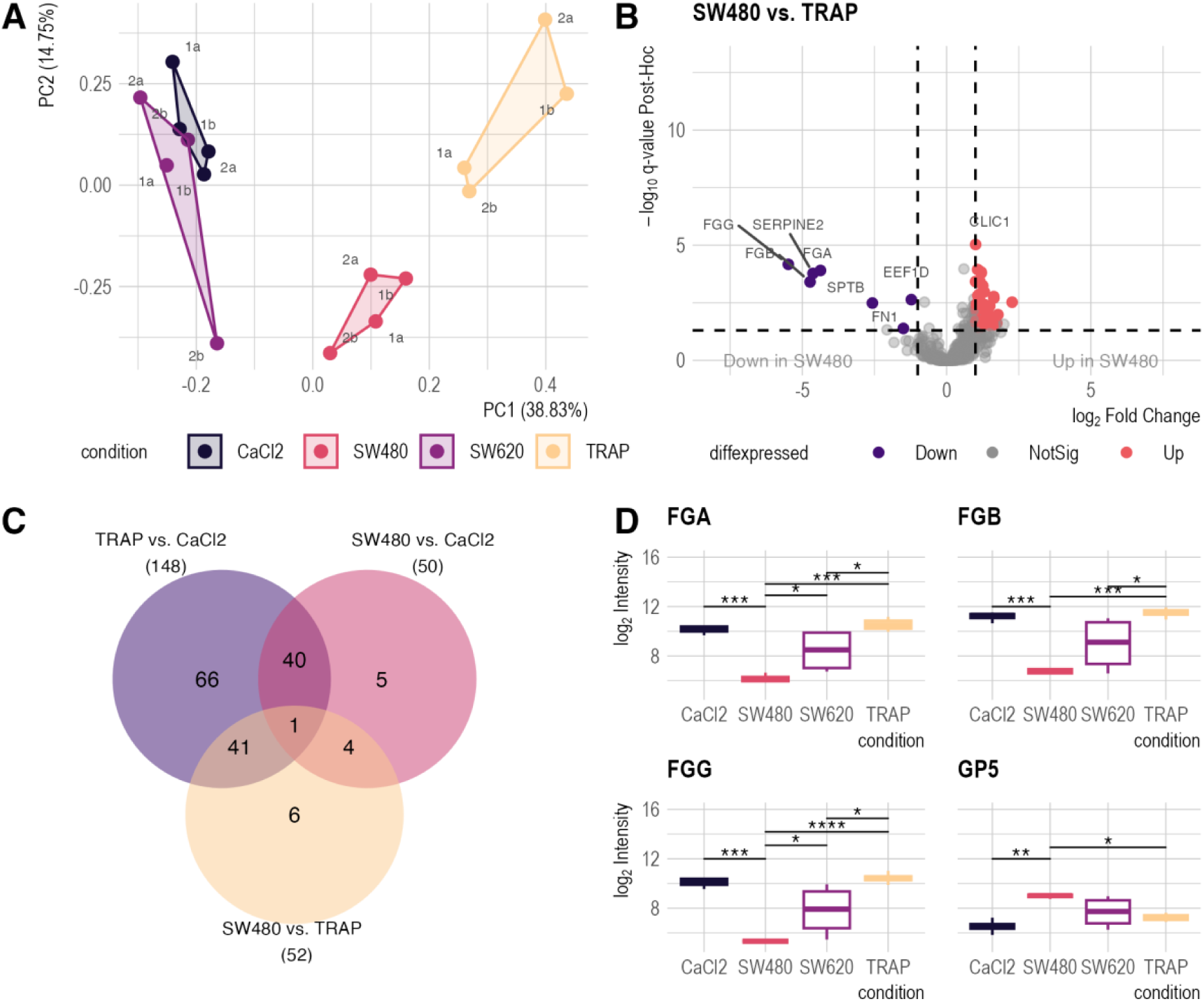
Proteomic analysis of the platelet secretome. **A:** Principal component analysis (PCA) plot of 546 secretome proteins. Clustering patterns mirror those observed in platelet sediment data (Figure 2A). PC1 accounts for 38.68% of the variance, and PC2 for 13.72%. **B:** Volcano plots illustrating differential protein expression in the secretome (ANOVA with Tukey HSD post-hoc, q ≤ 0.05, and |log_2_FC| of ≥1). Red points indicate proteins with significantly increased abundance, while purple points represent significantly decreased abundance of proteins. **C:** Venn diagram shows the set intersections of differentially abundant proteins identified in different conditions. Four proteins are uniquely differentially abundant in the secretomes of platelets activated by SW480 cells. **D:** Abundance patterns of four proteins specifically altered by SW480-induced platelet activation. Boxplots display protein levels across conditions. Asterisks indicate statistical significance (Tukey’s HSD post-hoc) q-value: * ≤ 0.05, ** q ≤ 0.05, *** q ≤ 0.001, **** q ≤ 0.0001.

### Changes within the tumor cell proteome are detectable after interaction with platelets

Using SILAC-based proteomics, we identified 650 unique tumor cell proteins. Notably, PCA exhibited a clear separation of both post-TCIPA SW480 and SW620 cells from their respective negative controls (Figure 5A, Supplementary Table S5). Furthermore, the cell lines exhibited divergent proteomes following TCIPA, suggesting distinct interaction modes with healthy platelets. A subsequent comparison of platelet-exposed SW480 cells with their untreated counterparts revealed a significant increase of 20 intracellular proteins and a significant decrease in the abundance of three proteins (Figure 5C, Supplementary Table S5, Additional File 1). Similarly, comparison of platelet-exposed SW620 cells with their naïve controls showed 20 proteins to be more abundant and 11 to be less abundant in the sediment (Figure 5D, Supplementary Table S5, Additional File 1). In addition to the comparison of platelet-exposed SW480 and SW620 cells to their naïve counterparts, we also analyzed the differentially expressed proteins between post-TCIPA SW480 to SW620 cells. It became apparent that the interaction with platelets led to distinct responses at the protein level: A total of 46 proteins exhibited differential abundance, with 26 showing decreased and 20 exhibiting increased abundance in SW480 cells compared to SW620 cells (Figure 5B, Supplementary Table S5, Additional File 1). To control for inherent differences between the cell lines, subsequent analysis focused on differentially expressed proteins uniquely identified in post-TCIPA SW480 and SW620 cells under naïve conditions. This approach guaranteed the elimination of initial differences between cell lines, highlighting the impact of platelet exposure. Twelve proteins exhibited significant abundance changes in both comparisons (naïve and platelet-exposed cells), while 251 proteins were exclusively different in naïve cells. These two sets of proteins were excluded from further analyses. The remaining 34 proteins, demonstrating unique changes in platelet-exposed tumor cells (Figure 6A & B), were selected for further investigations.

**Figure 5:**
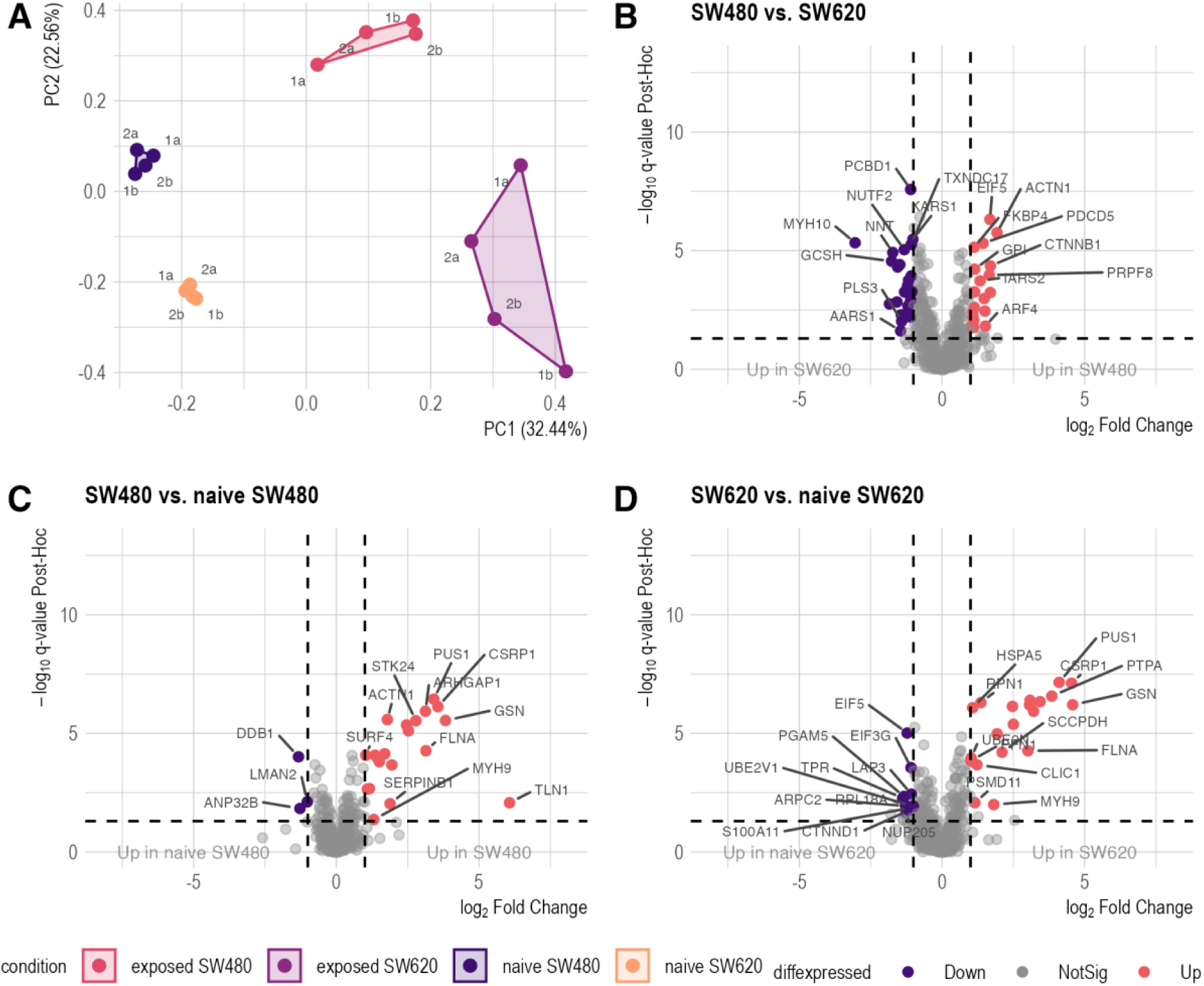
Analysis of tumor cell proteome changes upon platelet interaction. A: Principal component analysis of 650 proteins. Four distinct clusters are observed, with platelet-exposed SW480 and SW620 cells clearly separated from their respective controls (naïve cells). PC1 accounts for 32.44% of the variance, and PC2 for 22.56%. B-D Volcano plots showing differential expression analysis results (ANOVA with Tukey’s HSD post-hoc, q ≤ 0.05, and |log_2_FC| of ≥1. Red points indicate proteins with significantly increased abundance, while purple points represent significantly decreased abundance of proteins.

**Figure 6:**
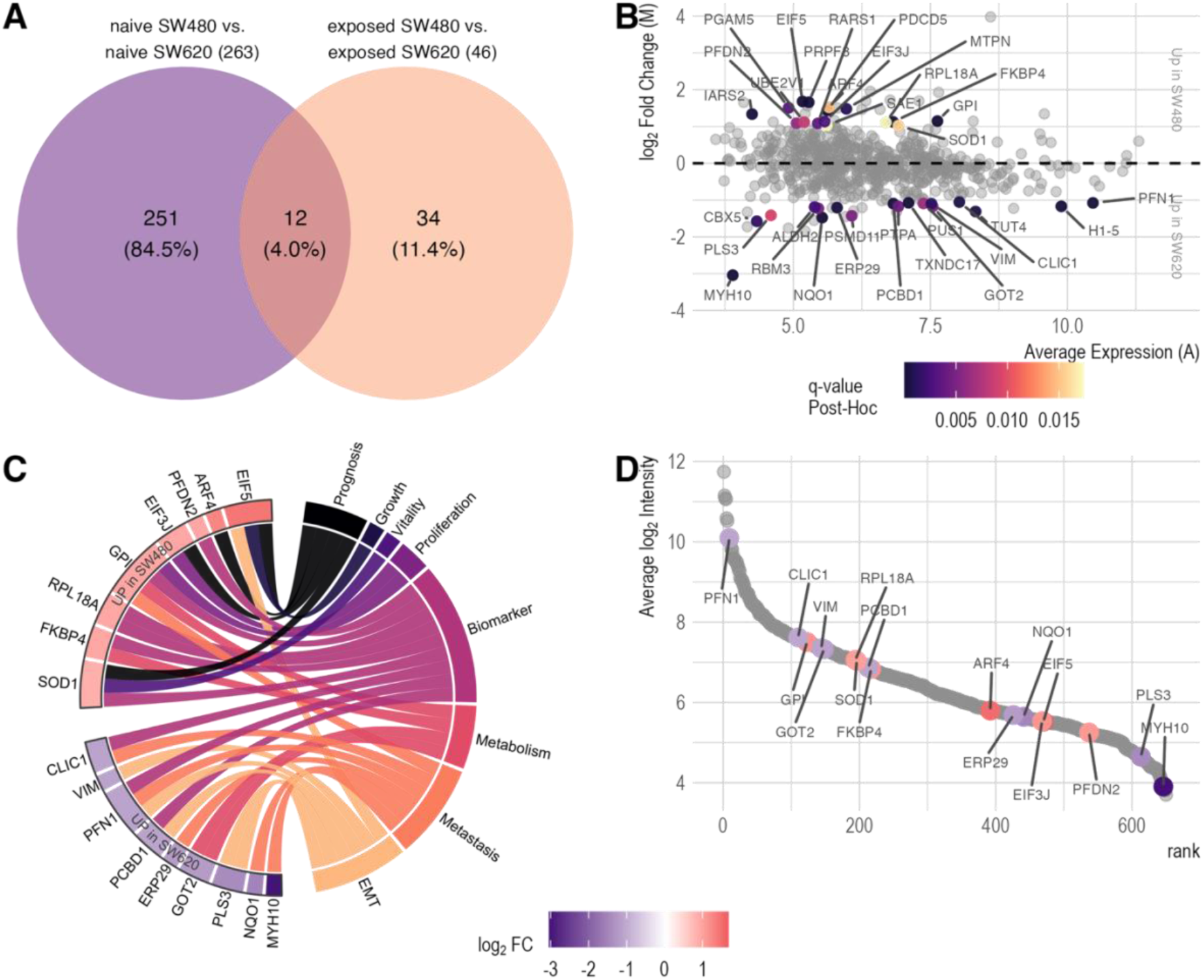
Identification and characterization of target proteins reveals association to cancer. **A:** Venn diagram showing the overlap of differentially expressed proteins between the cell lines before and after platelet exposure. 34 proteins are uniquely differentially expressed post-platelet exposure, correcting for baseline differences. **B:** Bland-Altmann/MA-plot visualizes the average expression of the tumor sediment proteins (A, x-axis) and the log_2_ fold change (M, y-axis) between platelet-exposed SW480 and SW620 cells (Figure 5D). The 34 TCIPA-unique proteins are highlighted, with color indicating the post-hoc q-value. **C:** Circular plot mapping the of 17 target proteins to colorectal cancer functions. Gene nodes are colored by log_2_FC between platelet-exposed SW480 and SW620 cells. **D:** Protein rank plot of 17 cancer/metastasis associated target proteins. Average log_2_ intensity (y-axis) was calculated across all naive and platelet exposed samples. Genes colored by log_2_FC between platelet exposed SW480 and SW620.

### A total of 17 target proteins were functionally associated with cancer

Overrepresentation analysis (ORA) was performed on the 34 identified targets using the MSigDB C2 collection. This analysis revealed 67 significantly enriched gene sets (q-value ≤ 0.05). Of these, eight gene sets contain the term ‘cancer’ and/or ‘metastasis’ in their descriptions, thus indicating the involvement of the 34 target proteins in cancerous diseases (Supplementary Table S6, Additional File 1 and Supplemental figure 4, Additional File 2). Seventeen of the 34 target proteins mapped to these eight cancer and metastasis-related gene sets. To further investigate their role in colorectal cancer, a literature review was conducted on these 17 proteins. Our review revealed a highly diverse involvement, with distinct patterns for proteins upregulated in platelet-exposed SW480 compared to SW620 cells. The eight target proteins, which showed increased abundance in platelet-exposed SW480 cells, have been previously described as prognostic or general biomarkers for CRC, promoters of tumor cell proliferation and growth, and factors associated with metabolism (Figure 6C, Supplementary Table S7, Additional File 1). Conversely, the remaining nine target proteins showed increased expression in platelet-exposed SW620 cells and have been implicated in the promotion of EMT and metastasis and play a role in cancer cell metabolism. The 17 target proteins encompass the full dynamic range of the protein quantification assay with a spread of over six orders of magnitude (Figure 6D).

## Discussion

Cancer metastasis remains a critical challenge in oncology, with hematogenous dissemination representing the primary mechanism of tumor progression. Platelets have emerged as crucial mediators in this process (6, 40): Platelets interact with circulating tumor cells (CTCs), shielding them from immune surveillance and providing essential survival factors. A key aspect of this interaction is tumor cell-induced platelet aggregation (TCIPA), a phenomenon observed across a range of cancer cell lines, including those derived from pancreatic and lung cancers (41–44). The formation of hetero-aggregates not only shields the tumor cells from shear stress and immune cell-mediated elimination but also supplies nutrients that support tumor cell survival, proliferation, and metastatic colonization (45–47). In this context, this study introduces a novel mass spectrometry-based analysis of TCIPA using a paired cell culture model consisting of a primary colorectal cancer cell line (SW480) and its lymph node metastasis derivative (SW620).

Proteomic analysis of SW480 and SW620 under naïve conditions revealed 263 differentially expressed proteins. The SW480 proteomic profile showed significant associations with EMT, adhesion, coagulation, motility and migration, and IL-2-STAT5 signaling. In contrast, the metastatic SW620 cells exhibited significant enrichment for MYC targets and increased MYC signaling. These findings strongly support the hypothesis that primary tumor cells undergo molecular reprogramming to facilitate detachment from the tumor mass, invasion and migration through the surrounding tissue, and finally entry into the circulation for metastatic dissemination. The upregulation of STAT signaling in SW480 cells aligns with Halim et al.’s (48) observations demonstrating its pro-tumorigenic effects, including inhibiting apoptosis, increased cell proliferation, migration, invasion, and dysregulating immune surveillance. The enrichment of MYC targets in SW620 cells underscores the importance of MYC in diverse oncogenic processes such as proliferation, apoptosis, differentiation, and tumor metabolism (49). While c-MYC overexpression has been implicated in mesenchymal-epithelial transition (MET) in certain cellular contexts (50), the precise role of MYC remains complex with studies associating MYC targeting with EMT (51, 52).

The analysis of primary tumor cells (SW480) and their paired lymph node metastasis cells (SW620) revealed significant differences in TCIPA capacity. Specifically, SW480 induced rapid, robust platelet aggregation, which was similar to the positive control (TRAP-6). In contrast, SW620 showed only weak and delayed interactions. This reduced TCIPA in metastatic SW620 cells correlates with their advanced metastatic stage, suggesting diminished reliance on platelet interactions after establishing lymph node metastasis. Proteomic analysis of SW480-stimulated platelet secretomes revealed decreased fibrinogen subunits (FGA, FGB, FGG) and increased Glycoprotein 5 (GP5), indicating a dual mechanism: sustained platelet adhesion via GP5-VWF interactions for tumor cell protection (53, 54), coupled with reduced fibrin clot formation to prevent entrapment. These findings demonstrate how tumor cells dynamically adapt platelet modulation strategies during metastasis progression, balancing circulatory survival with efficient dissemination.

In accordance with the presented results, it was most striking that SW620 cells exhibited substantial alterations in their own proteome following platelet interaction, although showing limited aggregation in LTA measurements and minimal impact on platelet proteome and secretome composition. This finding underscores the intricate nature of tumor cell-platelet interactions, extending beyond visible aggregation. Comparative analysis of platelet-exposed SW620 and SW480 cells revealed nine proteins with increased protein levels in the metastatic cell line, with four are associated with EMT (PLS3, VIM, PCBD1, PFN1) (55–69). All proteins have been observed as potential markers in CRC associated with tumor progression, prognosis and therapy resistance, e.g. via the influence of metabolic pathways or metabolic reprogramming (70–76). All these findings suggest that platelet exposure fosters a pro-metastatic, pro-EMT, and pro-metabolic environment in SW620 cells.

In contrast, the upregulation of several proteins in platelet-exposed SW480 cells, including EIF3J, EIF5, GPI, FKBP4, SOD1, PFDN2, SOD1, and RPL18A, suggests a multifaceted response that promotes colorectal cancer progression and metastasis through metabolic reprogramming (77), inflammatory processes (78, 79), and enhanced cell survival (80–84). Some proteins have already been described as prognostic biomarkers for CRC, implementing their relevance for CRC progression (85–93). These findings highlight the potential role of platelet interactions in driving a more aggressive CRC phenotype for the primary cell line and could suggest new avenues for therapeutic intervention.

## Conclusion

This study offers new proteomic perspectives on the complex relationship between colorectal cancer cells and platelets during metastasis. After the establishment of a novel approach to evaluate TCIPA, our findings demonstrate differential proteomic signatures and aggregation potential in primary tumor cells (SW480) and metastatic cells (SW620). While SW480 cells demonstrated robust platelet aggregation including proteome alterations associated with proliferation and metabolism, SW620 cells displayed attenuated aggregation but showed upregulation of EMT-related proteins. These results highlight the dynamic nature of tumor-cell platelet interactions throughout the metastatic cascade, underscore the potential importance of these interactions in promoting cancer progression, and establish a basis for future investigations into targeted anti-metastatic treatments.

## List of abbreviations

CRC: colorectal cancer
ANOVA: analysis of variance
CEP: cancer educated platelet
CTC: circulating tumor cells
EMT: epithelial-mesenchymal transition
FC: fold change
FDR: false discovery rate
GO: gene ontology
GSEA: gene set enrichment analysis
HSD: honestly significance difference
LC-MS/MS: liquid chromatography coupled tandem mass spectrometry
LTA: light transmission aggregometry
MAR: missing at random
MET: mesenchymal-epithelial transition
MNAR: missing not at random
MS/MS: tandem mass spectra
MSigDB: Molecular Signatures Database
ORA: overrepresentation analysis
PAR-1: preotease-actiated receptor 1
PCA: principal component analysis
PGI2: prostacyclin
PRP: platelet-rich plasma
SILAC: stable isotope labeling with amino acids
SWATH: sequential window acquisition of all theoretical mass spectra
TCIPA: tumor cell-induced platelet aggregation
TRAP(−6): thrombin receptor activator peptide 6

## Additional files

**Additional_File_1.xlsx:** Supplementary Table 1: Differential expression analysis results, naïve cell line proteome profile comparison; Supplementary Table 2: Selected Gene set enrichment analysis (GSEA) results for naïve cell line proteome profile comparison; Supplementary Table 3: Differential expression analysis results of platelet sediment proteome profile comparison; Supplementary Table 4: Differential expression analysis results of platelet secretome proteome profile comparison; Supplementary Table 5: Differential expression analysis results of cell line proteome profile comparison after TCIPA; Supplementary Table 6: Target proteine overrepresentation analysis results on MSigDB C2; Supplementary Table 7: Literature Review results on target proteins function in colorectal cancer and metastasis.

**Additional_File_2.docx:** Supplementary Figure 1: Immunofluorescence microscopy of CaCl_2_-exposed platelets; Supplementary Figure 2: Additional volcano plots of platelet sediment proteomics; Supplementary Figure 3: Additonal volcano plots of platelet secretome proteomics; Supplementary Figure 4: Functional analysis results of cancer and metastasis-associated genes using MsigDB C2.

## Declarations

### Ethics approval and consent to participate

The studies were approved by the local ethics committees of the University of Lübeck, namely #19-147-A. All patients provided informed written consent.

### Consent for publication

Not applicable.

### Availability of data and materials

The mass spectrometry proteomics data have been deposited to the ProteomeXchange Consortium via the PRIDE partner repository with the dataset identifier PXD062043.

### Authors’ contribution

**TS:** Conceptualization (supporting), Data curation (lead), Formal analysis (lead), Methodology (lead), Software (lead), Visualization (lead), and Writing – Original Draft Preparation (lead). **CG:** Writing – Original Draft Preparation (supporting). **KK:** Conceptualization (supporting) and Writing – Original Draft Preparation (supporting). **AR:** Investigation (equal). **LT:** Investigation (equal). **KS:** Investigation (equal). **JO:** Writing – Review & Editing. **LC:** Resources (supporting). **MK:** Methodology (supporting), Software (supporting). **AV:** Resources and Writing – Review & Editing (supporting). **TG:** Conceptualization (lead), Funding Acquisition (lead), Project Administration (lead), Resources (lead), Supervision (lead), Writing – Review & Editing (lead). All authors have agreed to the final version of the manuscript.

## Acknowledgements

We thank Julia Horn, Katja Klempt-Gießing, and Emma Neumann for their excellent technical assistance. Thorben Sauer gratefully acknowledges the scholarship he received from the Ad Infinitum Foundation. This work was supported by the German Network for Bioinformatics Infrastructure—de.NBI, service center BioInfra.Prot, funded by the German Federal Ministry of Education and Research (BMBF)—Grant FKZ 031 A 534A.

