## Additional File 2 for "Proteomic Remodeling During Tumor Cell-Induced Platelet Aggregation Unveils Metastatic Drivers in Colorectal Cancer"

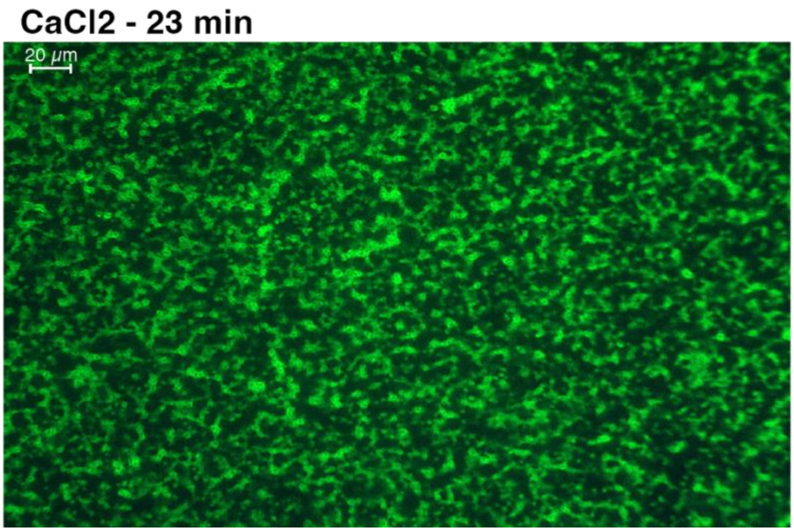


**Supplementary Figure 1:** Immunofluorescence microscopy of $CaCl_2$-exposed platelets. Platelets labeled with anti-CD42b antibody (green) after 23 minutes of $CaCl_2$ exposure. No visible platelet aggregation is observed, serving as a negative control for TCIPA experiments.

**Additional File 2** - Proteomic Remodeling During Tumor Cell-Induced Platelet Aggregation Unveils Metastatic Drivers in Colorectal Cancer – T. Sauer et al.

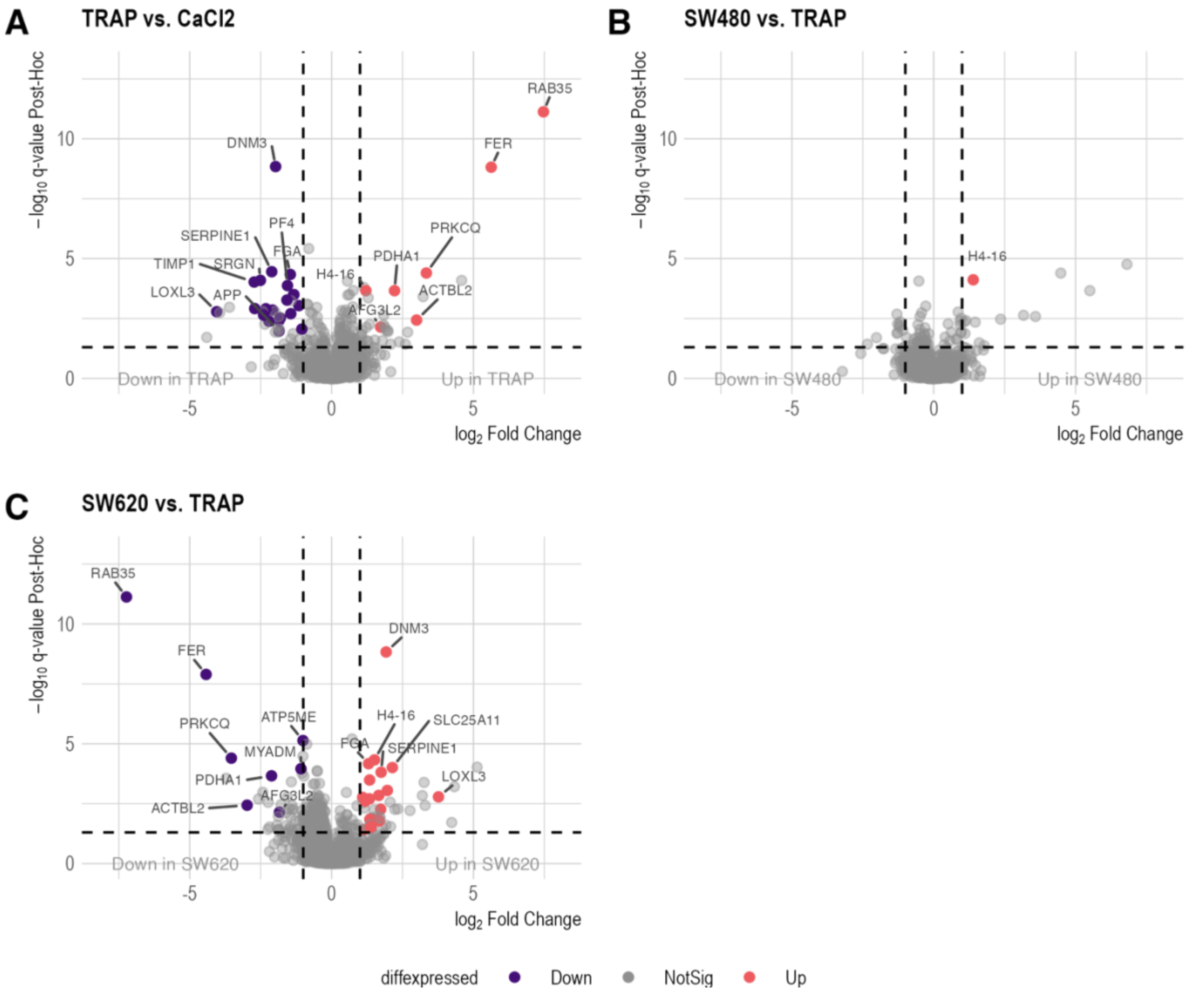


**Supplementary Figure 2:** Additional volcano plots of platelet sediment proteomics. A-D: Volcano plots illustrating differential expression in platelet sediments for three comparisons: A: TRAP *versus* CaCl2, B: SW480 *versus* TRAP and C: SW620 *versus* TRAP. Proteins are considered significantly abundant at ANOVA q-value and post-hoc q ≤ 0.05 and $|log_2FC|$ of ≥1. Significant proteins are highlighted in red, while downregulated proteins are shown in purple.

**Additional File 2** - Proteomic Remodeling During Tumor Cell-Induced Platelet Aggregation Unveils Metastatic Drivers in Colorectal Cancer – T. Sauer et al.

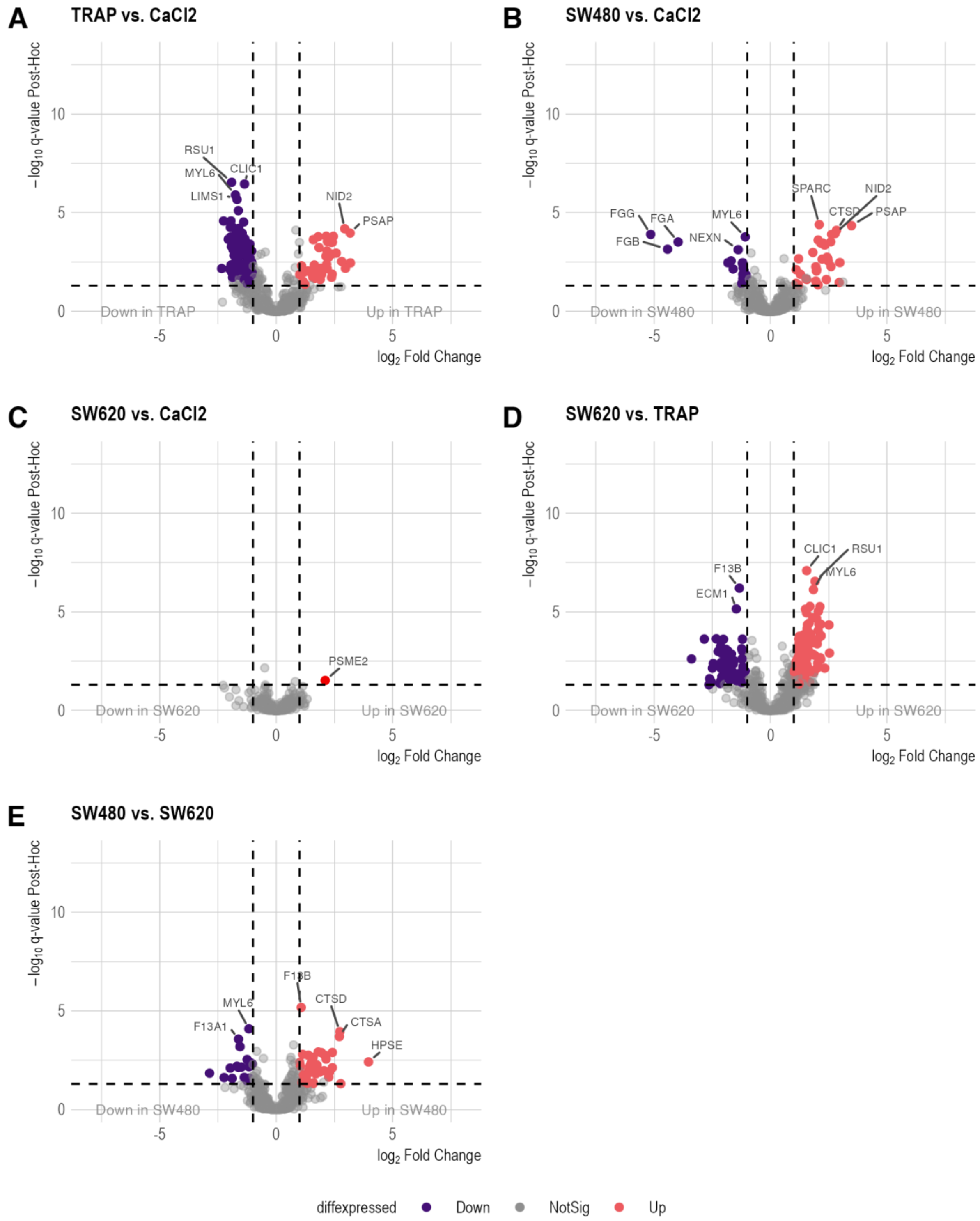


**Supplementary Figure 3:** Additional volcano plots of platelet secretome proteomics. A-E: Volcano plots illustrating differential expression in platelet sediments for three comparisons: A: TRAP *versus* CaCl2, B: SW480 *versus* $CaCl_2$, C: SW620 *versus* $CaCl_2$, D: SW620 *versus* TRAP, and E: SW480 *versus* SW620. Proteins are considered significantly abundant at ANOVA q-value and post-hoc q ≤ 0.05 and $|log_2FC|$ of ≥1. Significant proteins are highlighted in red, while downregulated proteins are shown in purple.

**Additional File 2** - Proteomic Remodeling During Tumor Cell-Induced Platelet Aggregation Unveils Metastatic Drivers in Colorectal Cancer – T. Sauer et al.

**Supplementary Figure 4:**

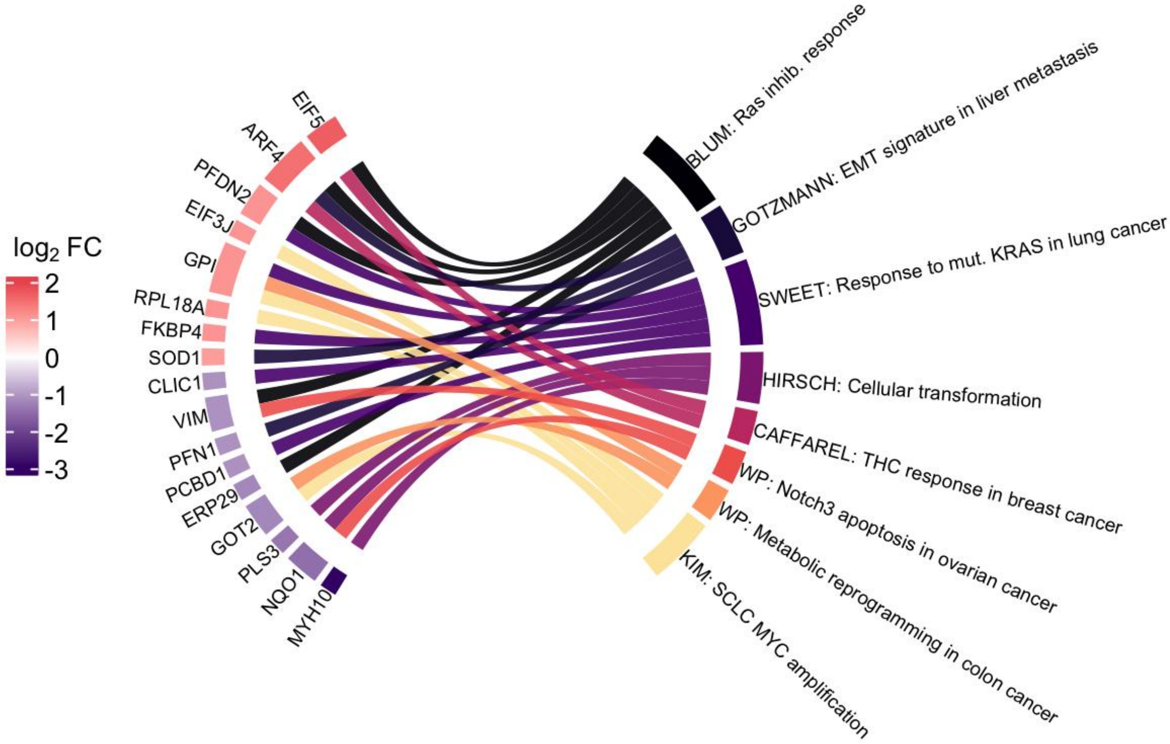


**Supplementary Figure 4:** Functional analysis of cancer and metastasis-associated genes. Chord diagram illustrating the results of an over-representation analysis using the MsigDB C2 collection. The diagram maps significantly enriched gene sets (q-value ≤ 0.05) containing 'cancer' and/or 'metastasis' terms to 17 cancer and metastasis-associated genes. Gene nodes are color-coded based on their log2FC between platelet-exposed SW480 and SW620 cells. Gene set labels display the author of the source publication (except for WP, denoting WikiPathways) and abbreviated gene set names for clarity.
